# *Twisted Sister1*: an agravitropic mutant of bread wheat (*Triticum aestivum*) with altered root and shoot architectures

**DOI:** 10.1101/2024.08.04.606531

**Authors:** Deying Zeng, Jiayu Peng, Lan Zhang, Mathew J. Hayden, Tina M. Rathjen, Bo Zhu, Zixian Zeng, Emmanuel Delhaize

## Abstract

We identified a mutant of hexaploid wheat (*Triticum aestivum*) with impaired responses to gravity. The mutant named *Twisted Sister1* (*TS1*) had agravitropic roots that were often twisted along with altered shoot phenotypes. Roots of *TS1* were insensitive of externally applied auxin with the genetics and physiology suggestive of a mutated *AUX/IAA* transcription factor gene. Hexaploid wheat possesses over eighty *AUX/IAA* genes and sequence information did not identify an obvious candidate. Bulked segregant analysis of an F_2_ population mapped the mutation to chromosome 5A and subsequent mapping located the mutation to a 41 Mbp region. RNA-seq identified the *TraesCS5A03G0149800* gene encoding a TaAUX/IAA protein to be mutated in the highly conserved domain II motif. We confirmed *TraesCS5A03G0149800* as underlying the mutant phenotype by generating transgenic *Arabidopsis thaliana*. Analysis of RNA-seq data suggested broad similarities between Arabidopsis and wheat for the role of *AUX/IAA* genes in gravity responses. Here we show that the sequenced wheat genome along with previous knowledge largely from the model species Arabidopsis, gene mapping, RNA-seq and expression in Arabidopsis have enabled cloning of a key wheat gene defining plant architecture.

## Introduction

Mutant analysis has been central for understanding fundamental processes in a diverse range of organisms that include microbes, plants and animals. In the plant kingdom, the identification and analysis of mutants have typically focussed on diploid species and preferably those with small genomes. This approach has enabled loss-of-function mutations to be more easily identified and cloned than in a polyploid species where homeologous genes can compensate for knock outs of single genes. In plants much of the research focus has been on small species with small genomes such as Arabidopsis (*Arabidopsis thaliana*) which has enabled the development of high throughput screening methods on Petri dishes or in small pots (Page and Grossniklaus, 2002). The study of roots provides added challenges as they are often hidden from view which further complicates mutant screening. Despite these difficulties, root mutants have been identified in crop species such as rice (*Oryza sativum*), barley (*Hordeum vulgare*) and maize (*Zea mays*), all of which have the advantage of being diploid (Parry et al., 2009). Rice is particularly attractive as its relatively small genome and abundance of genetic resources enables downstream activities such as the cloning of genes that underlie mutant phenotypes to be more easily undertaken than for species with larger genomes such as barley and maize (Viana et al., 2019).

Much research on species such as Arabidopsis and *Brachypodium distachylon* has been justified as being simple models for crop species despite the fundamental differences in genome structure. Wheat (*Triticum aestivum*) is one of humanities major crops providing about a fifth of the calories consumed by humans worldwide. It is grown in diverse locations and has been adapted to a wide range of conditions. The identification and characterisation of mutations in bread wheat is complicated by its hexaploid nature with its genome comprised of three related genomes (A, B and D). For instance, to find a recessive mutant commonly requires all three homeologs present in the three genomes of bread wheat to be knocked out to generate a phenotype. For example, recessive mutations in *ENHANCED GRAVITROPISM1* (*EGT1*) (Fusi et al., 2022) and *EGT2* (Kirschner et al., 2021) genes of barley confer a stronger root response to gravity than wild type resulting in steep roots. While single mutated genes in the diploid species barley conferred clear phenotypes, similar phenotypes in tetraploid durum wheat (*Triticum turgidum*) required knockout of homeolous genes on the A and B genomes. Furthermore, wheat has a large genome (∼17 Gbp) compared to other species such as rice (∼0.5 Gbp), maize (∼2.4 Gbp), barley (∼5 Gbp), sorghum (∼0.73 Gb) and Arabidopsis (∼0.12 Gb) (Page and Grossniklaus, 2002; Schreiber et al., 2018). Identification of a recessive mutation in a specific gene of Arabidopsis can be considered a relatively simple exercise if only a few thousand mutagenized seed need be screened on Petri dishes. By contrast, a mutation in an equivalent gene of wheat could require several hundred thousand seedlings to be screened if one simply considers the differences in genome size without accounting for gene redundancy due to the hexaploid genome. Furthermore, the large seed and plant sizes of wheat compared to Arabidopsis reduces the ability to devise high throughput screens particularly when searching for root mutants. Despite these disadvantages, Shorinola *et al*. (2019) identified a range of mutants in wheat with an altered number of seminal roots. Altering the morphology of roots in this way could provide advantages to root systems for the uptake of water and nutrients. The mutants were selected from a screen of targeting induced local lesions in genomes (TILLING) population of wheat but the genes underlying the mutants have not been identified. Zeng *et al*. (2024) recently described the *Stumpy* mutant of hexaploid wheat which is a conditional mutant with reduced cell elongation in roots when grown in high Ca^2+^ concentrations. A candidate gene of unknown function was identified and the absence of equivalent genes in Arabidopsis meant this species couldn’t be used as a convenient model system. Despite difficulties in using wheat as a model species, the availability of a sequenced genome, TILLING populations along with improved transformation methodologies go some way in compensating for these disadvantages. Although analysis of mutants in model species can suggest how equivalent mutated genes behave in wheat, it is still valuable to study the effect of mutations in the species of interest as a key step in translating research derived from model species to crops.

How plants sense and respond to a gravity stimulus has been elucidated largely by the study of root mutants in Arabidopsis. Plants sense gravity through specific cells (statocytes) which possess statoliths consisting of amyloplasts filled with starch granules (Nakamura et al., 2019). In roots it is the cells of the root cap that are the statocytes and these provide the sensory mechanism for detecting the direction of gravity. Under the influence of gravity, amyloplasts move in the direction of gravity to trigger a response that involves a change in the distribution of auxin such that roots bend towards the gravity vector. In shoot tissues, the response to gravity is opposite to roots such that shoots bend away from the gravity vector. Despite these opposing movements, the basic molecular processes appear to be shared by both roots and shoots (Sato et al., 2015; Nakamura et al., 2019).

For important crop species, the identification of gravity mutants and the cloning of underlying genes have been successfully undertaken in rice which is diploid and has a relatively small genome and barley with its larger genome but equivalent mutants in wheat are yet to be described in the literature. Here we describe the *Twisted Sister1* (*TS1*) mutant of hexaploid wheat. Growth of *TS1* under a range of conditions resulted in twisted or curled roots compared to the wildtype indicating that the response to gravity was perturbed. The mutation is semi-dominant and fine-mapping along with a transcriptomic analysis identified a mutation in a critical region of motif II of an *AUXIN/INDOLE-3-ACETIC ACID* (*AUX/IAA*) gene. These findings provide insight into the function of a wheat gene central in determining plant architecture in response to gravity.

## Results

### Isolation of *Twisted Sister1* and genetic analysis

From a screen using small pots of soil, we identified a seedling with seminal roots that emerged above the soil surface, a behaviour not seen in wild type (WT) seedlings (Fig. 1). Progeny of this plant confirmed the seedling as a mutant that segregated for the phenotype with mutant seedlings showing a similar behaviour when grown in agar medium (Fig. 1B). The mutant was named *TS1* in view of the twisted nature of roots. When grown in soil, progeny segregated into three groups based on shoot phenotypes. Seedlings were classed as WT, heterozygous *TS1* or homozygous *TS1* depending on the size of plants, number of stems and angle that stems made from the vertical orientation (Fig. 1C-E, Fig. S1). Severely-stunted seedlings with a single stem could be grown to maturity and formed a small head that developed from none to ten grains (Fig. 1). Grain from these plants all produced seedlings with the same phenotype which suggested the plants to be homozygous mutants. This was subsequently confirmed by maintaining *TS1* mutant lines, all with the same severe phenotypes, across five generations. By contrast, plants classed as heterozygous produced more seed than homozygotes and progeny that segregated confirming the parental plants as heterozygous. An agravitropic phenotype was apparent in roots of *TS1* plants from early germination with an altered shoot phenotype in heterozygous plants that persisted through to maturity (Fig. S2).

**Figure 1.**
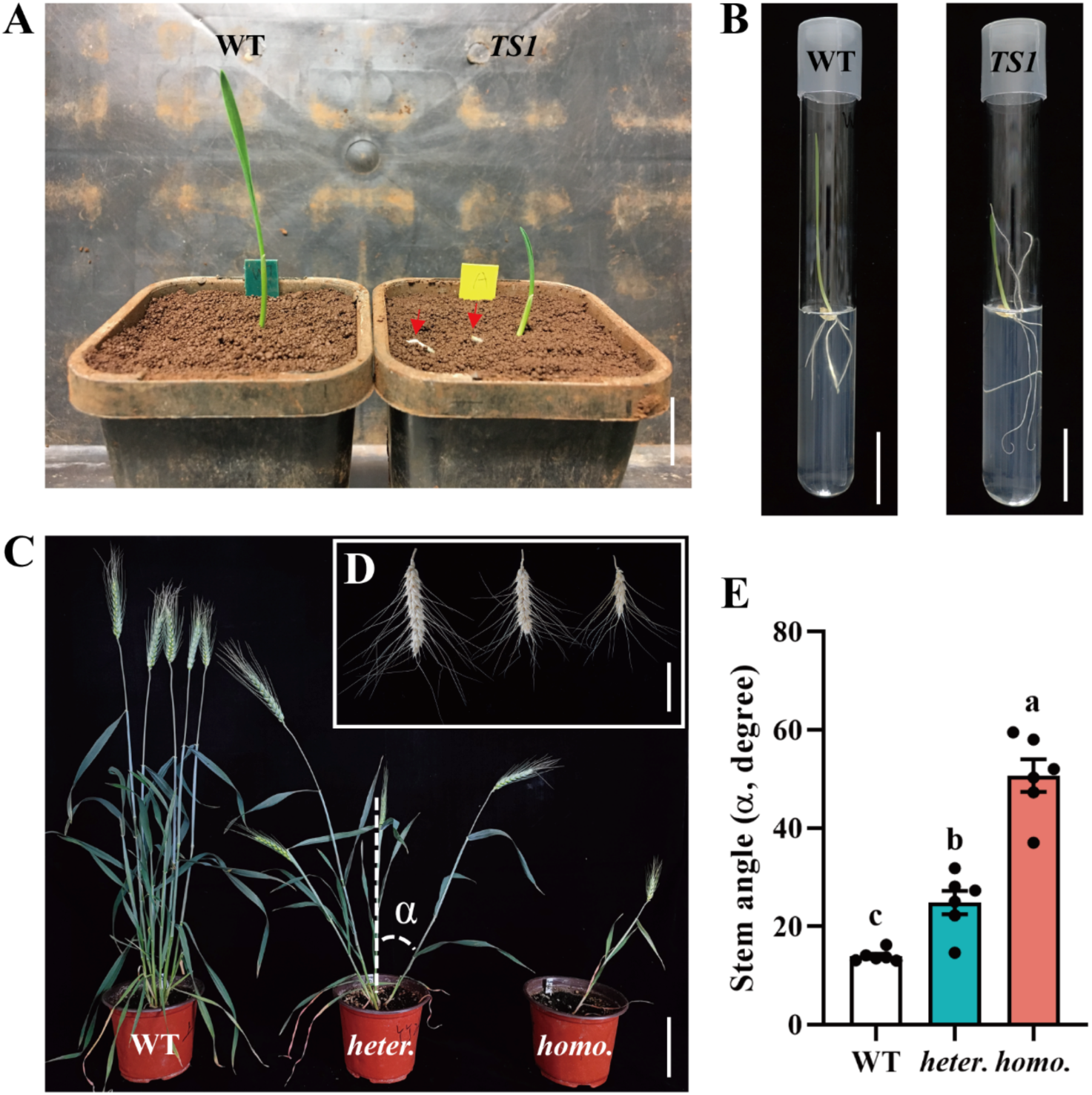
Root and shoot morphologies of WT and *TS1* mutant. **(A)** Wildtype and *TS1* seedlings grown in soil for 3 d (scale bar 2 cm). Red arrows indicate roots emerging from the soil surface. **(B)** Wildtype and *TS1* seedlings grown on nutrient agar under sterile conditions for 6 d (scale bar 3 cm). (**C)** Shoot phenotypes of WT, heterozygous *TS1* (*heter.*) and homozygous (*homo.*) *TS1* plants at maturity (scale bar 10 cm). **(D)** Dried mature heads of WT (left), heterozygous *TS1* (middle) and homozygous *TS1* (right; scale bar 5 cm). (**E**) Stem angles of WT, heterozygous (*heter.*) and homozygous (*homo.*) *TS1* plants at maturity. Stem angles (α) were measured as the angle subtended from the stem to the vertical upward direction as shown in **(C)**. Values shown are means ± SEM (n = 6). Different letters represent a significant difference at *P* < 0.05 as determined by a two-way ANOVA (Fisher LSD method, alpha = 0.05).

When grown in hydroponic culture, lines derived from heterozygous plants also segregated into three groups (Fig. 2). One group of seedlings had roots typical of WT phenotype, a group that was typified by twisty roots with few laterals and a group of small seedlings that along with the single stem, often produced only one or two seminal roots (Fig. 2). Seedlings classed as homozygous had more severe phenotypes than heterozygotes producing few short seminal roots that lacked lateral roots (Fig. 2D-G). Heterozygous *TS1* mutants developed the same number of seminal roots as WT with the primary seminal root longer than WT (Fig. 2D). However total root length of heterozygous *TS1* was considerably less than WT (Fig. 2G) on account of the fewer lateral roots (Fig. 2F). Internal cellular structure of *TS1* roots was also altered with the most notable difference being an absence of a pith which was evident in WT (Fig. 2H). The root phenotypes of *TS1* mutants varied (Fig S3) with some heterozygous seedlings forming tight twists in root sections (Fig. S3B, C, F) whereas other seedling had roots with larger bends not apparent on WT roots.

**Figure 2.**
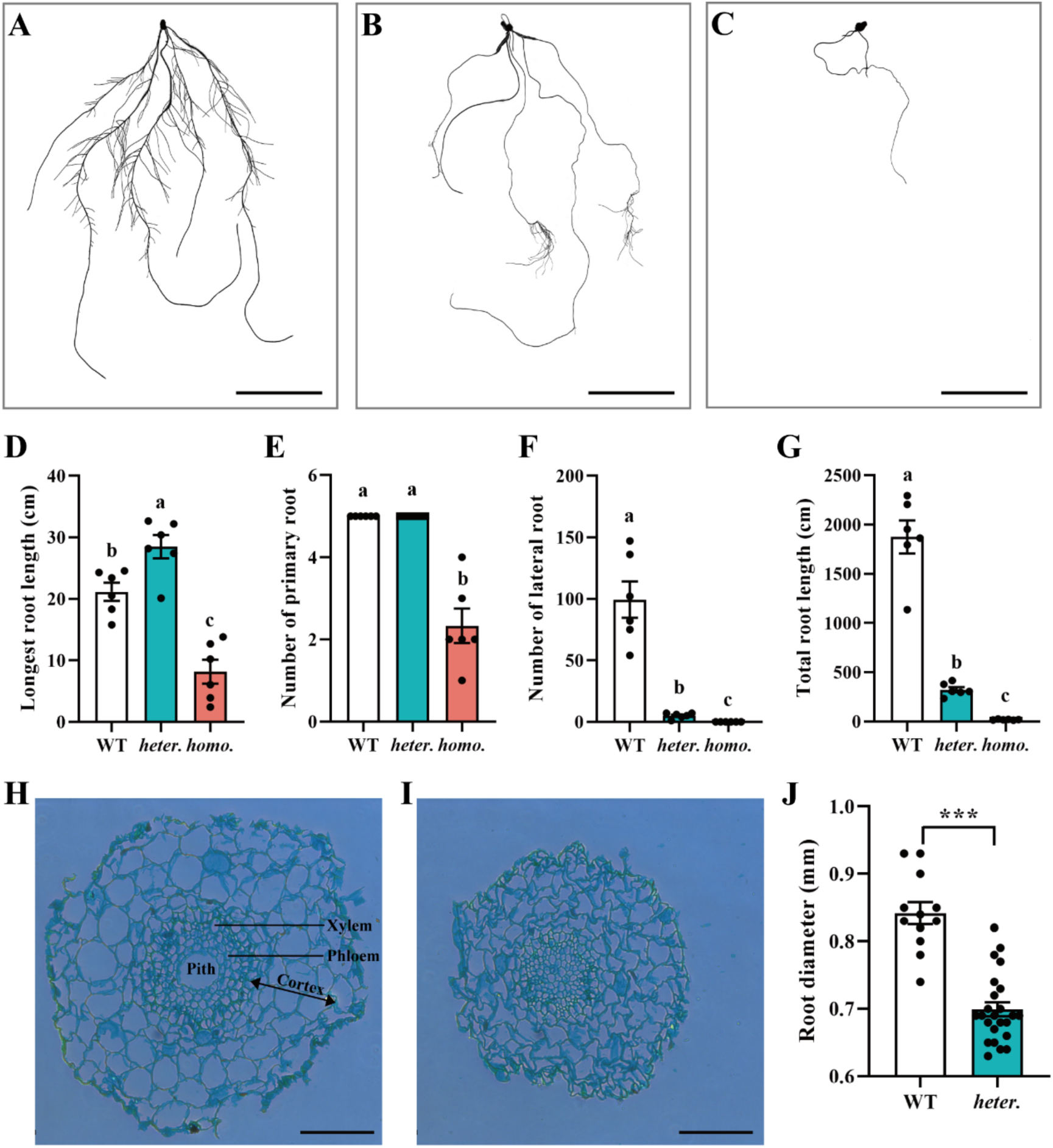
*TS1* mutants have altered root architecture. Root scans of **(A)** WT, **(B)** heterozygous *TS1* and **(C)** homozygous *TS1* lines grown for 9 d in hydroponic culture (scale bars = 5 cm). Nine-d old seedlings measured for **(D)** length of the longest seminal root, **(E)** number of seminal roots, **(F)** number of lateral roots and after 4 weeks growth **(G)** total root length. For (**D-E)**, values are means ± SE (n = 6). (**H**) Cross-sections of WT and (**I**) heterozygous *TS1* roots. (**J**) Root diameters of WT (n = 12) and heterozygous *TS1* (n = 24), values are means ± SE. Different letters represent a significant difference at *P* < 0.05 determined by a two-way ANOVA (Fisher LSD method, alpha = 0.05). Statistically significant differences between WT and *TS1* was determined with Student’s *t*-test: ***, P < 0.001.

To explore the genetic basis of the agravitropic phenotype of mutant *TS1*, WT cultivar (cv) Westonia and WT cv Baxter were each crossed with *TS1* to generate two segregating populations. F1 plants from both segregating populations had similar phenotypes to the parental line of mutant *TS1* having agravitropic roots and impaired development in shoots with large stem angles. Scoring the segregating population for shoot and root phenotypes as either WT, agravitropic or severely-agravitropic showed they segregated in 1:2:1 ratio that was not significantly different from ratios expected for a single semi-dominant gene controlling the agravitropic phenotype (χ^2^ = 1.14, *P* = 0.565 in 252 plants of the cross between *TS1* and Westonia; ξ^2^ = 0.43, *P* = 0.807 in 112 plants of the cross between *TS1* and Baxter).

### *TS1* mutants are insensitive of externally applied auxin and have altered accumulation of starch granules

Since growth of homozygous mutants was severely compromised and provided small amounts of grain, we chose to compare the physiology of *TS1* heterozygotes to WT. Mutant seedlings could be selected amongst segregating populations shortly after germination by observing roots and small homozygous *TS1* seedlings could then be removed prior to experiments. To quantify root responses to gravity, we grew seedlings in clear plastic pots inserted in black pots that enabled roots to be viewed and analysed when grown against the side of the plastic pot. For *TS1,* a proportion of seminal roots grew against the gravity vector while remaining roots either grew sidewards or downwards (Fig. 3A-C). By contrast, none of the WT roots grew upwards with all roots growing below the soil surface. In addition to the differences in root angles, *TS1* seedlings had shorter coleoptiles, larger angles between leaves and a smaller angle between the main stem and the horizontal position than WT (Fig. 3D-G). These phenotypes were indicative of perturbed gravity responses in both roots and shoots of *TS1*.

**Figure 3.**
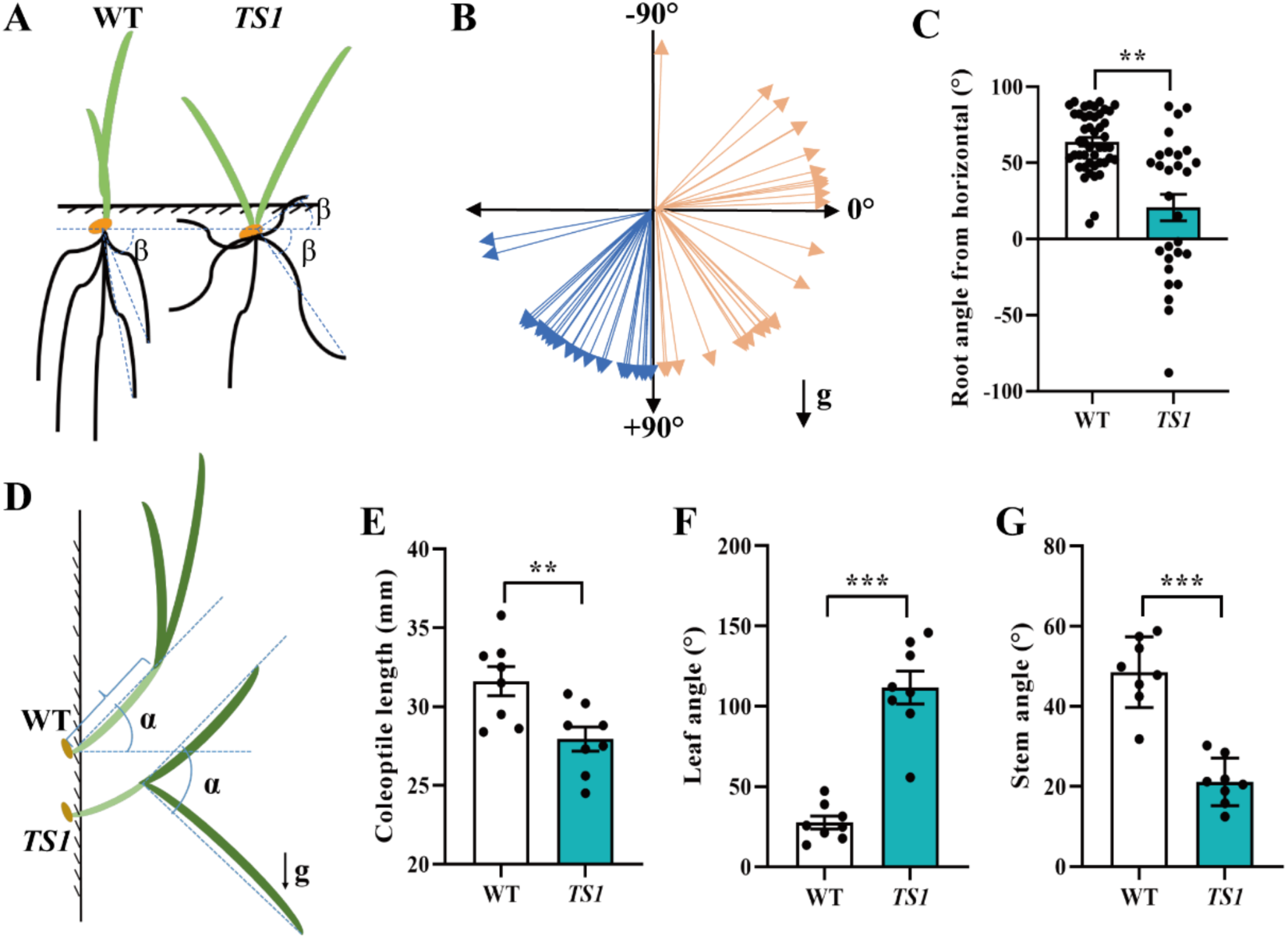
*TS1* roots and shoots are perturbed in their response to gravity. (**A**) Root angle (β) as measured by the angle subtended between the line from the root apex to the seed and the horizontal line (dotted lines). The horizontal position was taken as a reference and represented 0 degrees with a positive angle indicating roots that grew downwards while negative values indicated roots that grew upwards. **(B)** Vector diagram of root angles for *TS1* (orange) and WT (blue) plants. **(C)** Data of root angles plotted for WT (black bar) and *TS1* (turquoise bar) values are means ± SE (n = 28). **(D)** A diagram showing how coleoptile length, leaf angle and stem angle were measured for plants grown in pots for 14 d with g indicating the direction of gravity. **(E)** Coleoptile length, **(F)** leaf angle and **(G)** stem angle of WT (black bar) and *TS1* (turquoise bar) plants. All data were collected after 14 d growth and statistically significant differences between WT and *TS1* was determined with Student’s *t*-test: **, *P* < 0.01; ***, *P* < 0.001 (n = 8 for **E**-**F**).

Since auxin plays key roles in transducing the signal from gravity perception to changes in growth of plant tissues (Berleth et al., 2004; Leyser, 2018), we investigated the response of roots to externally-applied auxin. Typically, roots respond to auxin in a biphasic fashion with stimulation of root growth at low concentrations (less than 1 nM) whereas root growth is inhibited at higher concentrations (Leyser, 2018). When the auxin analog 1-naphthaleneacetic acid (NAA) was added to hydroponic solution, the longest root of *TS1* was insensitive of NAA at concentrations up to 0.1 μM with the longest root inhibited by about 27 % at 1 μM NAA (Fig. 4). By contrast, WT was inhibited at 0.01 μM with roots progressively inhibited at higher NAA concentrations (Fig. 4A-E). This finding showed that *TS1* is relatively insensitive of external auxin treatment compared to WT suggesting that the mutation in *TS1* had disrupted the auxin signalling pathway.

**Figure 4.**
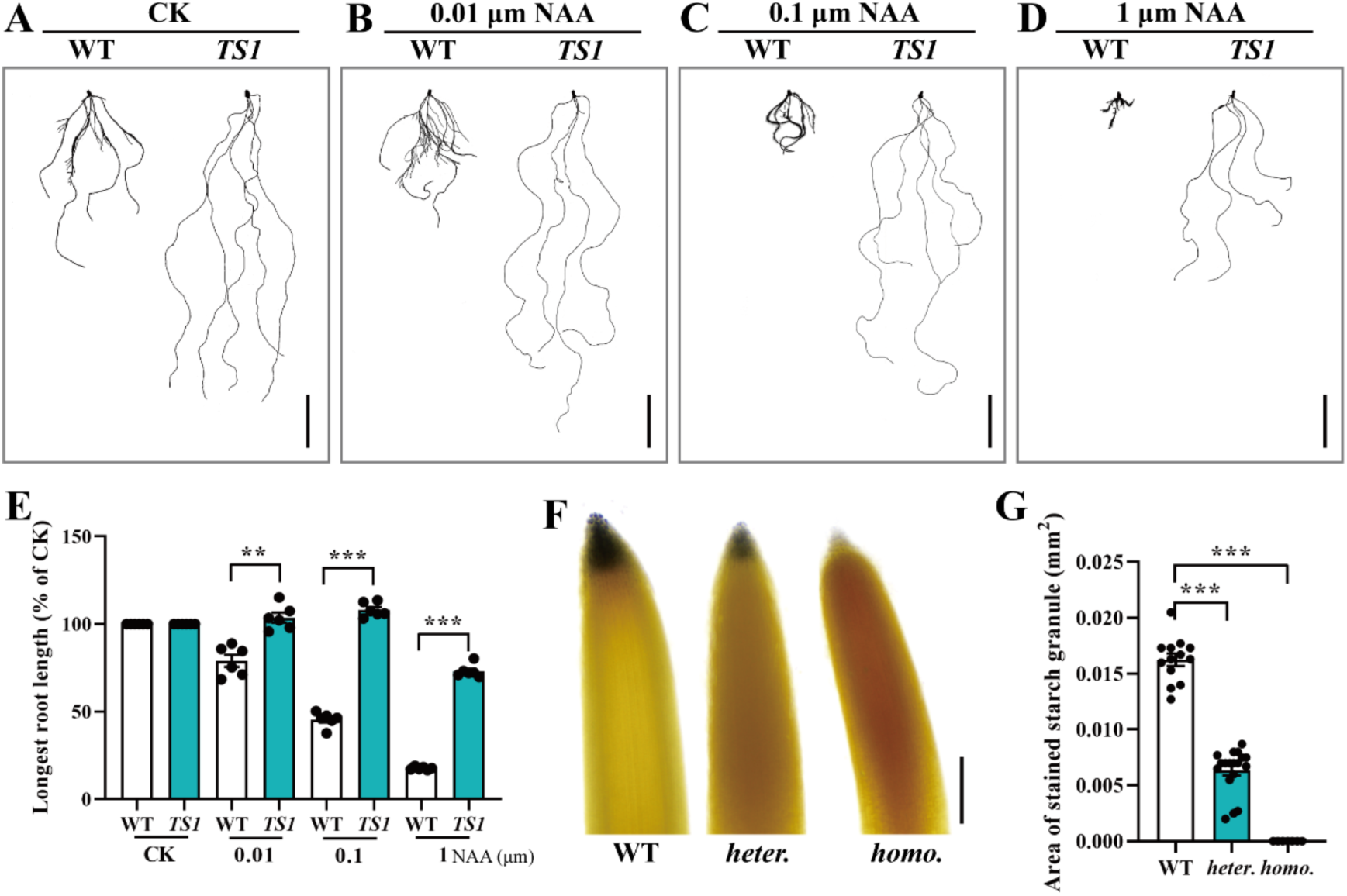
*TS1* is relatively insensitive of auxin and has reduced accumulation of starch granules. Root scans of WT and *TS1* grown in hydroponic culture for 9 d with solutions that contained **(A)** 0, **(B)** 0.01 μM, **(C)** 0.1 μM and **(D)** 1 μM of the auxin analogue NAA (scale bars = 5 cm). (**E**) The longest root length expressed as a percent of the control with no added NAA (error bars represent ± SE of 6 replicates). (**F**) Accumulation of starch granules in root tips of WT, heterozygous *TS1* (heter.) and homozygous *TS1* (homo.) plants grown in hydroponic culture without auxin treatment (scale bar = 0.5 mm). **(G)** The area of stained starch granules from the primary root of each line was measured for WT, heterozygous *TS1* (*heter.*) and homozygous *TS1* (*homo.*) plants (error bars represent ± SE of 12 replicates). Statistically significant differences (Student’s t-test): **, *P* < 0.01; ***, *P* < 0.001.

The agravitropic behaviour of *TS1* could have been due to the absence of starch granules in statocytes (root cap cells) as part of the early step of gravity perception. Although starch granules were totally absent from statocytes of homozygous *TS1* roots, statocytes of *TS1* heterozygotes still possessed starch granules albeit reduced in staining intensity compared to WT (Fig. 4F, 4G).

### *TS1* maps to chromosome 5A

To map the genetic location of *TS1*, we crossed *TS1* in a cv Westonia genetic background to WT in a cv Baxter genetic background. Segregant seedlings from the F2 population could be scored as having a WT or *TS1* phenotype so that bulked seedlings could be prepared for each phenotype. At the early seedling stage, the homozygous *TS1* seedlings could not be differentiated from heterozygous *TS1* seedlings so the mutant bulk consisted of a mixture of heterozygous and homozygous seedlings. The F2 bulks along with parental lines cv Westonia and cv Baxter were genotyped with an Illumina wheat 90K single nucleotide polymorphism (SNP) array. In total, 106 SNPs were moderately- or strongly-linked to one of the DNA bulks with *TS1* most likely located on chromosomes 2B or 5A (Supplemental Data Set S1, Fig. 5). Kompetitive allele specific (KASP) markers designed to both locations verified that *TS1* mapped to chromosome 5A. Further fine mapping with KASP and cleaved amplified polymorphic sequence (CAPS) markers identified *TS1* to be located between markers *KASP-IWB30321* and *CAPS-IWB54065* on the short arm of chromosome 5A with a physical distance of 46.12 Mbp that possesses 304 high confidence genes (Fig. 5B, Supplemental Data set S1). Although additional recombinant plants were available within this mapped region, the absence of polymorphisms between the parental lines restricted further mapping.

**Figure 5.**
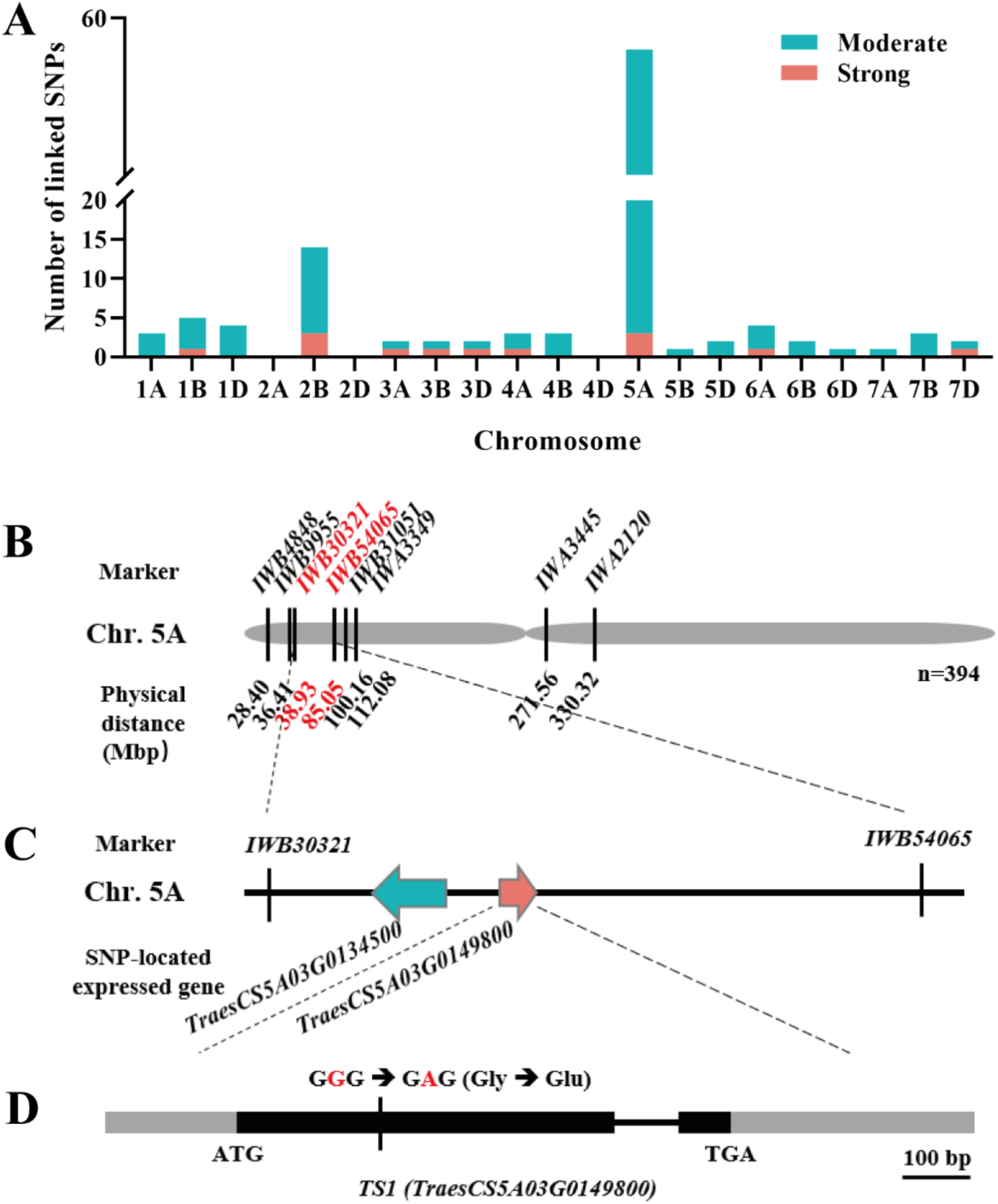
Cloning of *TS1* through a combination of mapping and transcriptomics. (**A**) Distribution of polymorphic SNPs moderately and strongly linked to the *TS1* locus on each wheat chromosome as identified by the 90K SNP array (Supplemental Data Set 1). (**B**) The *TS1* locus was mapped between the markers *IWB30321* and *IWB54065* on the short arm of chromosome (Chr.) 5A. (**C**) Transcriptomics identified two SNPs located in expressed genes in the region between markers *IWB30321* and *IWB54065.* RNA-seq data were analyzed to identify differentially expressed genes and SNPs within coding regions between WT and *TS1*. Shown are the orientations and order of the differentially expressed and annotated genes *TraesCS5A03G0134500* (turquoise) and *TraesCS5A03G0149800* (magenta). (**D**) Gene structure and sequence analysis of *TraesCS5A03G0149800* showing the location of the codon with the SNP (red) located within the coding region. The SNP changes the sequence from one encoding a glycine to one encoding a glutamate. The grey indicates untranslated regions while the black bar indicates the coding regions and the black line indicates an intron. Scale bar = 100 bp.

We then used an RNA-Seq analysis to search for the causative gene by exploring if expression level or a mutation within a coding region of a gene could be the basis of the mutant. A bioinformatics analysis of expressed genes in roots and shoots of heterozygous and homozygous *TS1* plants compared to WT identified 2 SNPs within genes located in the mapped region of chromosome 5A (genes *TraesCS5A03G0134500* and *TraesCS5A03G0149800*) (Fig. 5C). *TraesCS5A03G0149800* was considered a more likely candidate underlying *TS1* as it encodes a protein belonging to the *AUX/IAA* gene family whereas *TraesCS5A03G0134500* encodes a protein of unknown function. Sequence analysis of gene *TraesCS5A03G0149800* in mutant *TS1* showed that it had a SNP resulting in a single amino acid mutation changing a glycine to glutamate in the core domain II of the AUX/IAA protein (Fig. 5D). *TraesCS5A03G0134500* could be discounted as the causative gene since WT of the Chinese cultivar SM830 with normal gravity responses had the same sequence as *TS1.* Furthermore, F2 progeny of a cross between SM830 and *TS1* segregated for the *TS1* phenotype yet all plants had identical *TraesCS5A03G0134500* sequence regardless of phenotype (Fig. S4).

### *TS1* encodes an *AUX/IAA* gene

The mutation in the *AUX/IAA* gene of *TS1* was a strong candidate for conferring the agravitropic phenotype as mutations within the core domain II of *AUX/IAA* genes in other species such as Arabidopsis can result in dominant phenotypes (Luo et al., 2018). The *AUX/IAA* gene family in hexaploid wheat comprises over 80 members with *TraesCS5A03G0149800,* corresponding to the gene previously named as *TaIAA19* (Qiao et al., 2015), and here we have also adopted the shorthand designation *TaIAA19* for *TraesCS5A03G0149800*. *TaIAA19* was expressed throughout the WT cv Westonia plants with high expression in root and stem tissues and low expression in leaves (Fig. S5A). In addition to the SNP, relative *TaIAA19* expression in most tissues was reduced in *TS1* mutants compared to WT except in leaves and developing seed (Fig. S5B-F).

To verify that *TaIAA19* was the causative gene regulating root agravitropic and impaired growth phenotypes, we ectopically expressed both the WT and mutant versions of *TaIAA19* in Arabidopsis. None of the *AUX/IAA* genes of Arabidopsis are strongly related to *TraesCS5A03G0149800* with the protein encoded by the most similar gene sequence only having 38% similarity at the amino acid. Even though Arabidopsis is a dicot, it was effective in expressing *TaIAA19* with only the mutant version producing phenotypes of smaller shoots with petioles and leaves that curled or were slightly twisted (Fig. 6). Similarly, expression of only the mutant version of *TaIAA19* resulted in twisted and stunted roots typical of *TS1* (Fig. 6B-D). Importantly, expression of the WT version of *TaIAA19* did not generate phenotypes that differed from Arabidopsis WT plants. We assessed *TaIAA19* expression level to ensure that the phenotype caused by the mutated version of the gene was not simply due to greater expression even though expression in *TS1* wheat plants was often less than WT. Although different transgenic lines varied in *TaIAA19* expression level in roots (Fig. 6E) and shoots (Fig. 6F), the difference between WT and mutant versions of the gene could not be explained by expression level thus confirming the mutant version of the gene was responsible for generating the phenotypes. Other wheat proteins have related sequences to TaIAA19 and these, along with related sequences from Arabidopsis and rice were used to construct a neighbour-joining phylogenetic tree of AUX/IAA proteins (Fig. S6). In particular, rice OsIAA31 was the most similar AUX/IAA to TaIAA19 for which there is evidence for a role in roots (Fig. S6, Supplemental Data Set S2).

**Figure 6.**
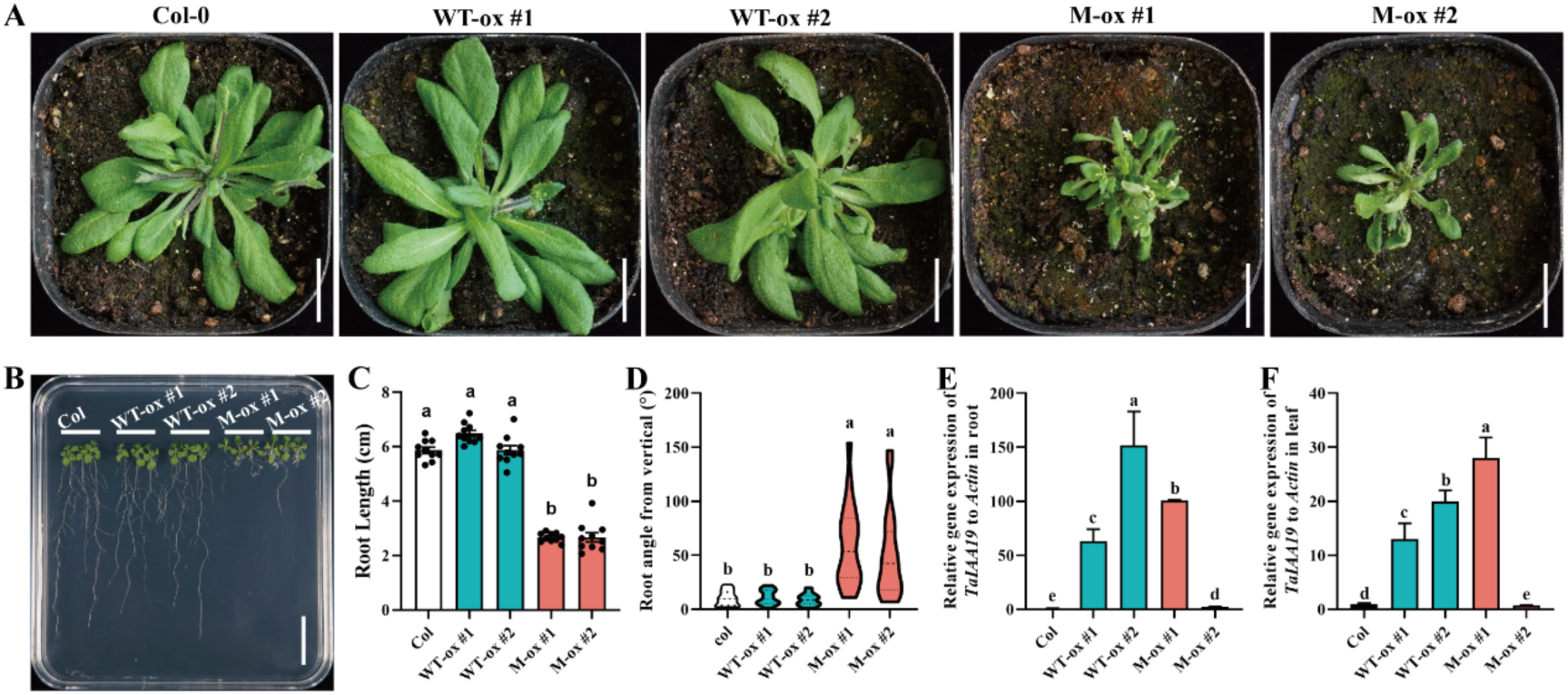
Functional confirmation of *TraesCS5A03G0149800* as the mutated gene underlying the *TS1* phenotype. (**A**) Shoot morphology of WT Arabidopsis (Col-0), Arabidopsis lines expressing the WT version of *TraesCS5A03G0149800* (WT-ox #1 and #2) and the *TS1* mutant version of *TraesCS5A03G0149800* (M-ox #1 and #2; scale bar = 2 cm). (**B**) One-week old seedlings of WT and transgenic lines expressing the WT and mutant versions of *TraesCS5A03G0149800* grown on a nutrient agar plate (scale bar = 2 cm). (**C**) Length of the longest root and **(D)** root angles from vertical of transgenic lines expressing the WT and mutant versions of *TraesCS5A03G0149800*. Seedlings were grown on nutrient agar plates for 7 d and values show means ± SE (n = 20). Level of *TraesCS5A03G0149800* expression in the Arabidopsis transgenics expressing the WT and *TS1* versions of *TraesCS5A03G0149800* shown relative to the internal actin gene in roots **(E)** and shoots **(F).** Error bars represent ± SE of 3 replicates. For all panels, different letters represent a significant difference at *P* < 0.05 as determined by a two-way ANOVA (Fisher LSD method, alpha=0.05). Scale bars indicate 2 cm in both panels **(A)** and (**B)**.

### RNA-Seq identifies *TaARF* and *TaPIN* genes regulated by *TaIAA19*

To identify genes regulated by *TaIAA19*, we undertook a bioinformatics analysis of the RNA-Seq data of the *TS1* lines and WT. The overall gene expression levels between three biological replicates in each genotype were highly correlated, with a correlation efficiency of > 0.86 (Pearson’s correlation; Supplemental Data Set S3). Homozygous *TS1* had thousands of genes differentially expressed when compared to WT in both roots and shoots with far fewer differentially expressed genes when heterozygous *TS1* was compared to WT (Fig. 7, Fig. S7; Supplemental Data Set S4). A total of 925 shoot (331 up-regulated and 594 down-regulated) and 428 root (129 up-regulated and 299 down-regulated) genes were differentially expressed when *TS1* mutants (heterozygous and homozygous combined) were compared to WT (Fig. 7A, Fig. S7; Supplemental Data Set S4 and S5). Gene Ontology (GO) analysis showed enrichment of these genes were related to auxin pathways (GO:0009734, GO:0009733), root development (GO:0010082 and GO:2000280) and cell wall biosynthesis (GO:0042546) (Figure 7B). The protein encoded by *TaIAA19* itself was classed in the auxin-activated signaling pathway (GO:0009734). We validated the differences in gene expression observed in the replicated RNA-Seq experiment by measuring gene expression of selected genes by qRT-PCR (Fig. 7 D-H).

**Figure 7.**
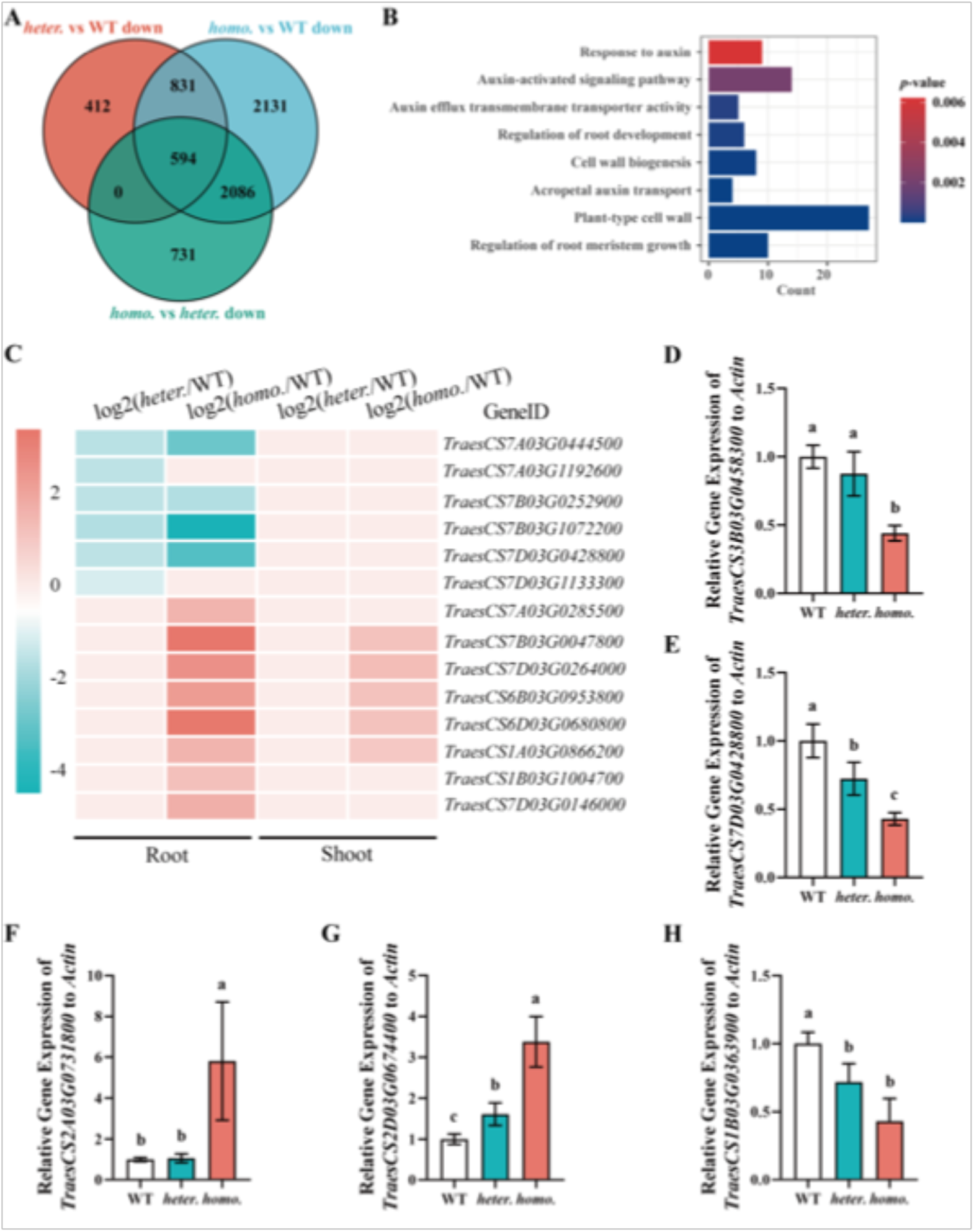
Analysis of WT and *TS1* transcriptomes of roots and shoots. (**A**) Venn diagram showing numbers of significantly downregulated root genes showing comparisons between heterozygous *TS1* (*heter.*) and WT (downregulated in *heter.*), homozygous *TS1* (*homo.*) and WT (downregulated in *homo.*) and, *homo. TS1* and *heter. TS1* (downregulated in *homo.*). (**B**) Enriched gene ontology terms for the 599 genes down-regulated in roots of both heterozygous and homozygous *TS1* seedlings. (**C**) Differentially expressed *PIN* (encoding directional auxin transport) and starch granule biosynthesis genes of *heter.* and *homo.* in the *TS1* mutant compared to WT. Gene identity (Gene ID) used the cv Chinese Spring v2.1 IWGSC nomenclature. The sequences of the proteins encoded by the first six genes (downregulated genes in *TS1* mutants) of the list cluster with sequences of Arabidopsis PIN proteins in a phylogenetic tree (Kumar *et al*. 2021). The upregulated genes encode proteins involved in starch biosynthesis (Supplemental Data Set 7). The heat map indicates genes whose expression is up-regulated (magenta) or down-regulated (turquoise) in *TS1* mutants compared to WT with the scale indicating approximate values as log2 transformed data. (**D-H**) Expression of selected genes in WT and *TS1* mutants as determined by qRT-PCR to validate RNA-seq data; *TraesCS3B03G0458300* (*TaARF9B*; Supplemental Data Set S3), *TraesCS7D03G0428800* (gene encoding a protein that clusters with Arabidopsis PIN proteins as shown by Kumar et al. 2021), *TraesCS2A03G0731800* (differentially expressed gene involved in starch biosynthesis GO:0019252), *TraesCS2D03G0674400* (differentially expressed gene involved in starch biosynthesis GO:0019252) and *TraesCS1B03G0363900* (differentially expressed gene involved in root development GO:0010082). Means (n = 3) of expression levels relative to a wheat actin gene are shown with different letters indicating a significant difference at *P* < 0.05 as determined by a two-way ANOVA (Fisher LSD method, alpha=0.05).

AUX/IAA proteins are involved in the early step of auxin perception and signal transduction and typically repress expression of specific auxin response factor (*ARF*) genes which themselves encode transcription factors (Cance et al., 2022). Previous studies on wheat *TaARF* genes have used conflicting nomenclatures and to avoid confusion we chose to use the gene nomenclatures of the cv Chinese Spring v2.1 International Wheat Genome Sequencing Consortium (IWGSC) along with the shorthand names originally provided by Qiao *et al*. (2018). To facilitate cross referencing, we provide Supplemental Data Set S6 which lists differentially expressed gene names originally designated by Qiao *et al*. (2018), the gene names designated by Jia *et al*. (2021) and both the version 1.1 and version 2.1 nomenclatures of the cv Chinese Spring IWGSC. The expression of one root and four shoot *TaARF* genes was down-regulated in the heterozygote compared to WT, while, expression of one root gene and three shoot *TaARF* genes was upregulated in *TS1* mutants (Supplemental Data Set S6). Since *PIN-FORMED* (*PIN*) genes and starch biosynthesis genes are implicated in gravity responses (Adamowski and Friml, 2015), we wondered if these genes were regulated by *TaIAA19*. The *PIN* gene family in hexaploid wheat comprises of 44 members (Kumar et al., 2021) and analysis of the RNA-Seq data identified 6 wheat genes belonging to two homeologous groups whose expression levels were downregulated only in roots of heterozygous *TS1* mutants compared to WT (Figure 7C), while none of the shoot *PIN* genes were affected. Twenty seven genes were upregulated in homozygous *TS1* roots but not in heterozygous *TS1* roots with 8 having similar sequence to Arabidopsis genes encoding proteins involved in starch biosynthesis (Supplemental Data Set S7). Genes with similar sequences to *PLASTIDIAL PHOSPHOGLUCOMUTASE* (*PGM*), *ADENOSINE DIPHOSPHATE GLUCOSE PYROPHOSPHORYLASE* (*ADG1*) and *STARCH SYNTHASE3* (*SS3*) all of which are regulated by auxin in Arabidopsis (Zhang et al., 2019), were upregulated in homozygous *TS1* mutants but were unchanged in the heterozygous *TS1* mutants compared to WT. Further analysis of genes involved in root development, root meristem development and cell wall biosynthesis showed that most genes were severely supressed or even silenced (FPKM<1), and displayed gene dosage effects in heterozygous and homozygous *TS1* mutants (Fig. 7B, Supplemental Data Set S4 and S5).

## Discussion

Here we identified the *TS1* mutant of bread wheat with altered responses to gravity. In particular, the roots behaved in an agravitropic fashion while the ability of stems and leaves to respond to the gravity vector was also altered. Growth of homozygous *TS1* seedlings was severely impaired, in which case we compared the physiology of heterozygotes to WT. The roots of *TS1* were insensitive of externally applied auxin and this provided clues as to the identity of the gene underlying the mutant. Mutations in *TaAUX/IAA* genes can confer dominant phenotypes with several resulting in roots with a relative insensitivity towards externally applied auxin (Luo et al., 2018). As noted in the Introduction, the identification of mutants, particularly root mutants, is a challenge in hexaploid wheat yet we identified *TS1* from a screen of only 2,000 seedlings. Key to the success in identifying *TS1* was its semi-dominance meaning the need for homeologs to also be mutated to generate a phenotype, as is often required for recessive mutants, could be avoided. Indeed, if other homeologs behave similarly to *TaIAA19*, then this would further reduce the number of seedlings needed for a screen to identify a mutant with an agravitropic behaviour.

We used a combination of gene mapping and transcriptomics to identify *TraesCS5A03G0149800* as the gene underlying the *TS1* mutant. By first mapping the mutation to within 46 Mb on chromosome 5A, we were than able to undertake a transcriptomic experiment to focus only on genes in this region that had mutations within coding regions. The agravitropic phenotype of *TS1* is semi-dominant which suggested a member of the wheat *TaAUX/IAA* gene family had been mutated based on published literature describing similar agravitropic phenotypes in other species, particularly Arabidopsis (Luo et al., 2018). However, there are few studies on functions of *TaAUX/IAA* genes in wheat with none focussing on gravity responses. The *TaAUX/IAA* family comprises of some 83 members (Qiao et al., 2015) such that identifying a candidate gene based simply on sequence information (DNA or protein) was not possible. Furthermore, although the *AUX/IAA* genes seemed to be obvious candidates, other genes when mutated can confer dominant phenotypes in plants some of which are associated with gravity responses (Naoi and Hashimoto, 2004; Meinke, 2013; Zhang et al., 2014; Marques-Bueno et al., 2021).

Dominant phenotypes in a range of plant species have been described with many due to mutations within a core domain of *AUX/IAA* genes. The proteins encoded by WT *AUX/IAA* typically act as repressors of gene expression and this repression is alleviated in the presence of auxin which results in degradation of AUX/IAA protein thus enabling transcription to occur (Luo et al., 2018). Mutations in the core domain II of AUX/IAA having the conserved amino acid sequence GWPPV stabilise the association with promoter regions and prevents transcription to occur. The specific mutation in *TS1* occurred within core domain II changing a Gly residue to a Glu residue. This same change occurred in the *OsIAA23* gene of rice encoding an AUX/IAA protein resulting in a semi-dominant mutant with a range of root phenotypes including an altered response to gravity (Jun et al., 2011). To confirm the SNP was responsible for the gravity phenotype in *TS1*, we expressed both the WT and mutant versions of *TaIAA19* in Arabidopsis. Arabidopsis expressing only the mutant form of the gene showed a stunted and twisted root phenotype. Because root phenotypes can occur due to over-expression of WT versions of *AUX/IAA* genes, we verified that expression level did not explain the difference between mutant and WT versions of the *TaAUX/IAA* gene when expressed in Arabidopsis. Although AUX/IAA proteins of Arabidopsis are distantly related to TaIAA19 protein (Figure S6), it’s apparent that TaIAA19 mutant protein shares sufficient similarity to Arabidopsis AUX/IAA proteins involved in gravity response to confer phenotypes that emulate the *TS1* mutants (twisted roots and small shoots; Fig. 6).

The transcriptomics experiments using RNA-seq identified many genes with perturbed expression in *TS1* mutants. Many of the genes with reduced expression are associated with root development (GO:0010082 and GO:2000280) and the observed altered root structures of *TS1* mutants, both internally and externally, are consistent with these data (Fig. 7A, B and Fig. 2). In particular, we focussed on *ARF* genes as early specific targets of repression by AUX/IAA proteins (Cance et al., 2022). As expected, the expression of several *TaARF* genes was repressed in heterozygous *TS1* compared to WT (Supplemental Data Set S6) consistent with the role of AUX/IAA as a transcriptional repressor resulting from increased stability of the mutant form of the protein. However, surprisingly the expression of a similar number of *TaARF* genes was increased in heterozygous *TS1* which appears to counter the idea that AUX/IAA proteins always act as transcriptional repressors. However, ARFs can undertake a large variety of transcriptional regulatory mechanisms and are not restricted to activator and repressor roles (Cance et al., 2022). Interestingly, the *TaMOR* gene that regulates crown root initiation in wheat was shown to act through a *PIN* gene (Li et al., 2022). From the transcriptomics data we identified expression of six *TaPIN* genes that was reduced in heterozygous *TS1* plants compared to WT (Fig. 7C and Supplemental Data Sets S4 and S5). This finding is consistent with *TaARF* genes acting as part of the signal transduction pathway in wheat to regulate gravity responses through *TaPIN* expression. *PIN* genes encode transporters of auxin and these transporters have been shown to be involved in the redistribution of auxin across root cells to initiate a change in root direction under the influence of gravity (Adamowski and Friml, 2015). Surprisingly, expression of genes involved in starch biosynthetic pathways were increased in *TS1* homozygotes which appears inconsistent with the absence of starch granules in root cap cells whereas the same genes were not perturbed in the *TS1* heterozygotes despite reduced staining for starch granules (Fig. 4F, Fig. 7C). In the loss-of-function mutant *pin2* of Arabidopsis, expressions of *PGM*, *ADG1* and *SS4*, genes involved in starch granule synthesis, were all upregulated resulting in increased starch granules accumulating in the root apex (Zhang et al., 2019). While an inverse relationship between expression of *PIN* genes (reduced) and genes involved in starch biosynthesis (increased) appear consistent between Arabidopsis and homozygous *TS1* mutants, they appear to differ markedly in how this affects accumulation of starch granules.

While the *TS1* mutant does not have direct application in improving the agronomic performance of bread wheat, it enabled us to identify a key gene controlling root and shoot architectures. Mutations in the conserved domain II of *TraesCS5A03G0149800* underlying *TS1* appear to generate severe phenotypes with no direct agronomic value. However, it is possible that mutations targeted to other regions of *TraesCS5A03G0149800* using new CRISPR/CAS methods (Ni et al., 2023) could confer more subtle phenotypes that are beneficial.

## Materials and Methods

### Germplasm

Seed of wheat cv Westonia was treated with 1 mM sodium azide using a previously described method (Chandler and Harding, 2013). Twenty M3 families were kept separated from one another to ensure that mutants identified from different families could be maintained as independently-derived mutants. A WT line of cv Westonia, was developed by single seed descent to ensure a consistent genetic background when backcrossing and generation of a mapping population.

### Soil screen

Germinated seed of the M3 generation of mutated cv Westonia lines were planted into small pots as described previously (Delhaize et al., 2015), except four seedlings were planted per pot. Growth of seedlings was monitored daily with attention paid to any roots that showed aberrant behaviours such as growing on the soil surface or into the air. After three to five days, soil and seedlings were tipped into a tray and roots viewed more closely to identify any potential mutants. About 2,000 seedlings were screened using this soil-based method. Selected seedlings with an aberrant behaviour were transferred to large pots for growth to maturity. Grain was collected and the progeny screened using the same method as the primary screen. Only plants that produced progeny showing a similar phenotype to the parental plant were confirmed as mutants. One seedling was confirmed to be a gravity-responding mutant with some roots that grew against gravity. This seedling was grown to maturity and crossed to WT cv Westonia as an initial step to generate backcrossed germplasm. The mutant was also crossed to cv Baxter, a different Australian wheat cultivar, to generate a mapping population. The F1 seedling of the mutant crossed to WT cv Westonia was further backcrossed up to five times to cv Westonia. Since the mutation initially appeared to be dominant, progeny seedlings with a phenotype could be selected with a screen using large Petri dishes as described below.

### Phenotyping

For physiological experiments *TS1*was backcrossed three times to WT cv Westonia. To phenotype roots, plants were grown in transparent plastic pots (ANOVA pots 20 cm diameter and 10 cm height; http://www.anovapot.com/php/anovapot.php) filled with potting mix that were placed into same-sized black pots as described by Richard *et al*. (2015). Plants were grown in a greenhouse set to 25°C. Roots of seedlings were analysed by marking the root tips and measuring the angle that a line drawn from the tip to the seed made in relation to a horizontal line. The clear pots were photographed when plants had reached maturity. The root angle of Arabidopsis transgenic lines was measured by the angle between the root apex to the seed and the vertical downward gravity direction when plants were grown on half-strength Murashige and Skoog medium in agar for one week.

Seedlings were grown by hydroponics using a solution at pH ∼ 5.9 with a composition described previously (Delhaize et al., 2004), except that 0.5 mM NH4NO3 was substituted with 0.5 mM NH4Cl and 0.5 mM KNO3. The biological buffer 2-(N-morpholino) ethanesulfonic acid hemi-sodium (MES) was added to 1 mM to stabilise the pH at 5.9. Germinated seedlings were placed in small, netted baskets inserted into the lid of a closed container (20 l) ensuring roots were submerged in the nutrient solution. Plants were grown in a cabinet with a daily cycle of 16 hours light at 23°C and 8 hours dark at 15°C with a light intensity of 740 µmol m^-2^ s^-1^. To assess the auxin sensitivity of root growth, the auxin NAA was added to the nutrient solution and root elongation measured. Twenty days after sowing, the length of the longest root was measured and the roots stored in 50-70% ethanol. Shoots were dried at 65°C and weighed. Harvested roots were rinsed in water and spread out on a transparent tray atop a flatbed scanner (Epson Expression 10000XL) after which roots were scanned at a resolution of 400 dpi and total root lengths and average root diameters analysed using WinRHIZO Pro v.2013 (Regent Instruments Inc., Quebec, QC, Canada). For shoots, the angle that stems made relative to the horizontal was measured for plants grown to maturity in pots. The angle made by the stem in relation to the horizontal position and the angle between the first two emerging leaves were measured with a protractor.

### Rapid seedling screen

Seedlings were germinated on moistened blotting paper inserted into Petri dishes and grown for several days. The *TS1* mutants could be scored as seedlings with roots that grew in a random fashion where they often “lifted” off the base of the dish and showed a degree of twisting or curling (see Fig. S2B). Roots of WT seedlings occasionally lifted off the Petri dish but typically remained relatively straight and did not have random growth. Seedlings that could not be scored with confidence over an initial 3 d were grown for a further period until their phenotype could be confidently scored. Since *TS1* roots had an agravitropic behaviour, it was expected by chance some would grow downwards and could be incorrectly scored as WT. Allowing seedlings to grow for a longer time reduced the likelihood of incorrect scoring as additional seminal roots emerged.

### Mapping of the *TS1* mutation

The *TS1* mutant was crossed to cv Baxter to generate a mapping population. Single nucleotide polymorphisms (SNPs) associated with the mutation were identified using an approach based on bulk segregant analysis (Michelmore et al., 1991). A pair of contrasting bulks was prepared by pooling equal amounts of genomic DNA from fifteen F2 seedlings from each group that were selected based on the phenotypes of *TS1* or WT. An artificial F1 sample was prepared by combining an equal amount of DNA from each of the parental lines. The bulked DNA samples, artificial F1, cv Baxter and *TS1* (cv Westonia background) were genotyped for 90,000 gene-based SNPs using the Infinium iSelect 90K wheat bead chip array (Wang et al., 2014), following the manufacturer’s instructions (Illumina Ltd). The SNPs were assessed for putative linkage by comparing the normalised theta values for each sample as described by Hyten *et al*. (2008). SNPs were considered to be putatively linked to the mutation when the normalised theta values for the *TS1* bulk and *TS1* parent (cv Westonia background, control line), and WT bulk and cv Baxter were similar, and the normalised theta value for the artificial F1 samples was about halfway between that of the other samples. Putatively linked markers were confirmed by manual inspection using GenomeStudio v2011.1 software (Illumina Ltd).

Polymorphic SNPs between cv Westonia and cv Baxter within identified genomic regions were used in KASP and CAPS assays to verify they co-segregated with the *TS1* trait and to further map the gene. The *TS1* trait initially appeared to be dominant so seedlings in the F2 of the *TS1* by Baxter cross with a WT phenotype were selected and SNPs identified as being polymorphic between parents were used in KASP and CAPS assays to map the mutation. The phenotypes of selected seedlings in the F2 were verified in the F3 generation. Sequence data (RefSeq v2.0) of cv Chinese Spring from the IWGSC was accessed to identify candidate genes within the region mapped with flanking markers (Alaux et al., 2018).

### Observation of starch granules and paraffin sectioning

To observe starch granules in the root tips, *TS1* mutant and WT seeds were germinated on moist petri dishes and root tips stained by a modification of a previously described method (Kitomi et al., 2020). Briefly, collected root tips (1.5 cm in length) from 3-d-old seedlings were dipped in 1% I2-KI solution for 45s. The stained roots were washed with distilled water, spread on glass slide, and then observed under a stereo microscope (Nikon SMZ745T). The stained area in the root tip was measured by ImageJ software (https://imagej.nih.gov/ij/). The protocols used to process root tip samples and to perform paraffin sectioning followed a previously described method (Cheng et al., 2018). Briefly, root tips which were excised as a section of 0.5-1.0 cm from the apex were fixed for 2 d at 4°C in a solution comprising of 2.5 % glacial acetic acid (v/v), 35 % (v/v) ethanol and 1.85 % formaldehyde (v/v) mixed in water. Samples were treated with a series of dehydration and infiltration steps and then embedded in the paraffin. Tissue sections (10 μm thick) were cut with a rotary microtome (Leica, Germany), treated with xylene and stained with 1% fast green (Sigma, Germany) and observed under a biological inverted microscope (Leica, Germany). Primary roots from at least 3 seedlings of each genotype were observed.

### Genetic Transformation of Arabidopsis

Full-length coding sequences of WT and mutant forms of candidate gene *TaIAA19* were cloned into the pCAMBIA1302 vector under the control of the 35S promoter. The plasmid was introduced into *Agrobacterium tumefaciens* strain GV3101 to transform Arabidopsis ecotype Columbia 0 (Col-0) using the floral dip method (Zhang et al., 2006). Progeny seedlings were analyzed for root angles using the method described above for wheat.

### Phylogenetic analysis

The amino acid sequences of AUX/IAA proteins in Arabidopsis (*Arabidopsis thaliana*), rice (*Oryza sativa*), and wheat (*Triticum aestivum*) were retrieved from EnsemblPlants (https://plants.ensembl.org/index.html). Sequence alignment of these AUX/IAA proteins was conducted using the MUSCLE programs. The phylogenetic tree was constructed using the Neighbor-Joining method by using the MEGA11 software (Tamura et al., 2021).

### RNA extraction, RNA-Seq and real-time quantitative PCR

Roots and leaf samples, which for *TS1* both showed mutated phenotypes, were used for RNA-seq. Seedlings were grown by hydroponics for two weeks as described above for phenotyping. Samples were harvested from individual plants, and each line had three biological replicates. The biological replicates of RNA-seq libraries were constructed and sequenced using an Illumina NovaSeq 6000 platform with the mode of 150 bp pair end (PE). All RNA-seq libraries in this study were developed and sequenced (Novagene, Beijing, China). Analysis of RNA-Seq data was conducted using a previously described method (Guo et al., 2023).

Various tissues from WT and *TS1* mutant grown in little pots were collected in liquid nitrogen. Total RNA was extracted from three biological replicates using an OminiPlant RNA Kit (CWBIO, Jiangsu, China) according to the manufacturer’s instructions, and first strand cDNA was synthesized using the EasyScript® All-in-One First-Strand cDNA Synthesis SuperMix Kit (TransGen Biotech, Beijing, China). The cDNA was diluted with water in a 1:5 ratio, and 2 μl of diluted cDNA was used for qRT-PCR with the 2xSuperFast Universal SYBR Master Mix (CWBIO) using a CFX96 Real-Time PCR Detection System (Bio-Rad Laboratories, Inc.). The wheat *β-ACTIN* gene was used as an endogenous control (Paolacci et al. 2009). Primers used for qRT-PCR are listed in Supplemental Data Set S8.

## Author contributions

LZ assisted with the phenotyping experiment, JP and ZZ undertook the RNA-Seq analysis, MJH arranged SNP chip analysis, TMR identified the mutant and undertook preliminary characterisation. DZ and ED led the project and undertook the bulk of genetic and physiological analysis of *TS1*. All authors contributed towards writing the manuscript.

